# Impact of cilia-related genes on mitochondrial dynamics during *Drosophila* spermatogenesis

**DOI:** 10.1101/2021.07.03.450990

**Authors:** Elisabeth Bauerly, Takuya Akiyama, Kexi Yi, Matthew C. Gibson

## Abstract

Spermatogenesis is a dynamic process of cellular differentiation that generates the mature spermatozoa required for reproduction. Errors that arise during this process can lead to sterility due to low sperm counts and malformed or immotile sperm. While is estimated that 1 out of 7 couples encounter infertility, the underlying cause of male infertility can only be identified in 50% of cases. Here, we describe and examine the genetic requirements for *missing minor mitochondria* (*mmm)*, *sterile affecting ciliogenesis* (*sac)*, and *testes of unusual size* (*tous)*, three previously uncharacterized genes that are predicted to be components of the flagellar axoneme. Using *Drosophila*, we demonstrate that these genes are essential for male fertility and that loss of *mmm, sac,* or *tous* results in complete immotility of the sperm flagellum. Cytological examination uncovered additional roles for *sac* and *tous* during cytokinesis and transmission electron microscopy of developing spermatids in *mmm, sac,* and *tous* mutant animals revealed defects associated with mitochondria and the accessory microtubules required for the proper elongation of the mitochondria and flagella during ciliogenesis. This study highlights the complex interactions of cilia-related proteins within the cell body and advances our understanding of male infertility by uncovering novel mitochondrial defects during spermatogenesis.

## Introduction

*Drosophila* has the longest sperm in the animal kingdom that has currently been identified. Species such as *Drosophila bifurca* reach nearly 6cm in length (1,000x the length of human sperm) (Pitnick et al., 1995). To reach these extreme lengths, some important modifications are needed during the spermatozoon flagellum formation. Two such modifications include the elongation of mitochondria to extend along the entire length of the axoneme and the formation of the accessory microtubules that bud off of the β-tubule of the axoneme (Kiefer, 1970). In humans and most other mammals, the flagellum does not require the additional mitochondrial support and only extends to about 1/3 the length of the axoneme. While little is known about accessory microtubules, they are involved in the elongation of the mitochondria that flank the axoneme (Fuller, 1993).

Spermatogenesis is the process by which single germline stem cells go through a series of divisions and complex cellular remodeling events to generate mature spermatozoa. In *Drosophila*, a germline stem cell divides to produce spermatogonia that go through 4 additional rounds of mitosis. Due to incomplete cytokinesis, these divisions result in a cyst that contains 16 cells that are connected through cytoplasmic bridges. These cells will complete the division process with two rounds of meiosis, resulting in 64 interconnected spermatids (Fuller, 1993, White-Cooper, 2003, Demarco et al., 2014). The final step in this developmental sequence is a remodeling phase referred to as spermiogenesis. During spermiogenesis, the flagellum is assembled concurrently with the remodeling of the nucleus and elongation of the mitochondria (Tokuyasu et al., 1972, Fabian & Brill, 2012). These processes all require the spatiotemporal coordination of hundreds of proteins that, if disrupted, can lead to sterility.

Dyneins are a class of microtubule motor proteins that are essential for nearly every stage of spermatogenesis. These multi-protein motor complexes are primarily involved in retrograde transport and are divided into two classes, cytoplasmic and axonemal. Cytoplasmic dyneins are involved in nuclear positioning, spindle formation and function during mitosis and meiosis, and the transportation of proteins and organelles both within the cell body and in the cilium (Palmer et al., 2009). Axonemal dyneins are the complexes that form the inner and outer arms along the axoneme, the core component of cilia, which provide the force required for sperm motility (Hou & Witman, 2015, Roberts et al., 2013). To investigate the regulation of ciliogenesis during spermatogenesis, we screened dynein-related proteins in *Drosophila* that were reported as having high expression levels in the testes (Chintapalli et al., 2007). From this screen, we identified 3 genes, *CG14838*, *CG14651*, and *CG4329*, that are predicted to be components of the axonemal dynein complex. Utilizing mutant alleles to interrogate the function of these genes, we discovered that they are essential for male fertility with critical functions within the cell body during spermatogenesis that impact mitochondrial dynamics during axonemal elongation.

## Results

### *CG14838, CG14651*, and *CG4329* encode dynein-related proteins

Based on sequence similarities, CG14838 is predicted to be a member of the intermediate chain of the axonemal dynein complex and a component of the inner dynein arm. It is projected to be the homolog of Dynein Axonemal Intermediate Chain 3 (DnaI3), mutations of which are affiliated with developmental abnormalities, including occipital encephalocele and brain lobulation, which can affect the formation of both the body and the brain (Hofmeister et al., 2018). While there is no prediction for the biological function of CG14651 in *Drosophila*, it is speculated to be orthologous to Dynein axonemal heavy chain 1 and 10 (DNAH1 and 10), suggesting it may be involved in axoneme biogenesis and the formation of the inner dynein arms (Oiwa & Sakakibara, 2005). *CG4329* is the homolog of *cilia and flagella associated protein 57 (CFAP57)*, which is reported to be involved in the assembly of the inner dynein arms in *Chlamydomonas* (Bustamante-Marin et al., 2020). Here, we characterize *CG14838*, *CG14651*, and *CG4329*, which we have named *missing minor mitochondria* (*mmm)*, *sterile affecting ciliogenesis* (*sac)*, and *testes of unusual size* (*tous),* respectively. Our data indicate that these genes are essential for spermatogenesis and have unexpected roles outside of ciliogenesis that impact cell division and mitochondrial dynamics that contribute to infertility.

### *mmm*, *sac*, and *tous* are required for male fertility

*Drosophila* Mmm, Sac, and Tous have been identified as putative dynein-related proteins through sequence similarities, and prior microarray analysis further reveals peak expression levels in the testes (Chintapalli et al., 2007). To determine the function of each gene in spermatogenesis, we obtained mutant alleles and males homozygous for each mutation were collected within 24 hours post-eclosion. Testes from each of the three genotypes were then surgically removed and examined for morphological abnormalities. Live imaging of the testes indicated that the overall morphology was unperturbed in animals homozygous for *mmm^ET1^* and *sac^ET1^* (Figure 1). However, testes from *tous^ET1^* homozygotes displayed a spectrum of morphological defects including a reduction in size and irregular shape of one or both testes (Figure 1G-H, and M). The homozygous males collected from each of the 3 genotypes were kept isolated from females for 2 weeks and examined again at the end of this time frame to ascertain if there were any age-related morphological defects or in sperm production. *mmm^ET1^* and *sac^ET1^* displayed no indication of morphological abnormalities and age did not appear to exacerbate the morphological irregularities observed in *tous^ET1^* (Figure 1A-H). The only other notable defect that arose during this time was the failure of mature sperm to be loaded into the seminal vesicles, an indication of defective spermatogenesis (*red* arrows, Figure1A-H and N). To investigate the capacity of these mutant animals to generate mature sperm, seminal vesicles and testes were dissected and analyzed using immunofluorescence in fixed tissue as well as live imaging. Examination of testes stained with anti-pan polyglycylated Tubulin antibodies revealed that all three mutants were capable of generating mature sperm (Figure 1I-L). However, live imaging revealed that the sperm failed to individualize or coil and were immotile, resulting in disorganized bundles of sperm (Figure 1I-L and Supplemental Movie 1). These data suggest that ciliogenesis is disrupted in these mutants and that all three fail to complete the last steps of spermatogenesis. Consistently, progeny counts from homozygous males mated to *wildtype* females confirmed that *mmm*, *sac*, and *tous* mutants are 100% male sterile (Figure 1O). We validated the mutant phenotype by complementation tests with a genomic deficiency line (*Df*). As expected, *mmm^ET1^/Df30580* and *sac^ET1^c/Df7620* displayed the same male-sterile phenotype observed in the homozygous animals (*mmm^ET1^/ Df30580, n*=10/10; *sac^ET1^/ Df7620, n*=10/10). These results indicate that they are likely null alleles. Unfortunately, we could not perform a fertility assay for *tous^ET1^/Df*282 males as they were not able to survive to adulthood, suggesting that the *tous* mutation is likely a hypomorph or that the combination of the *tous* mutation with the disruption of another gene in the deficiency line is toxic. To further confirm our findings, rescue constructs were generated for all three mutants that contained an mScarlet-I tag at their C-termini. We were able to rescue the male-sterile phenotype of *mmm* and *sac* mutants (*mmm^ET1^*, *n=*9/10; *sac^ET1^*, *n=*9/10). However, *tous* fertility was not restored by the rescue construct presumably due to interference from the tag rendering the protein non-functional (*n*=0/10).

**Figure 1.**
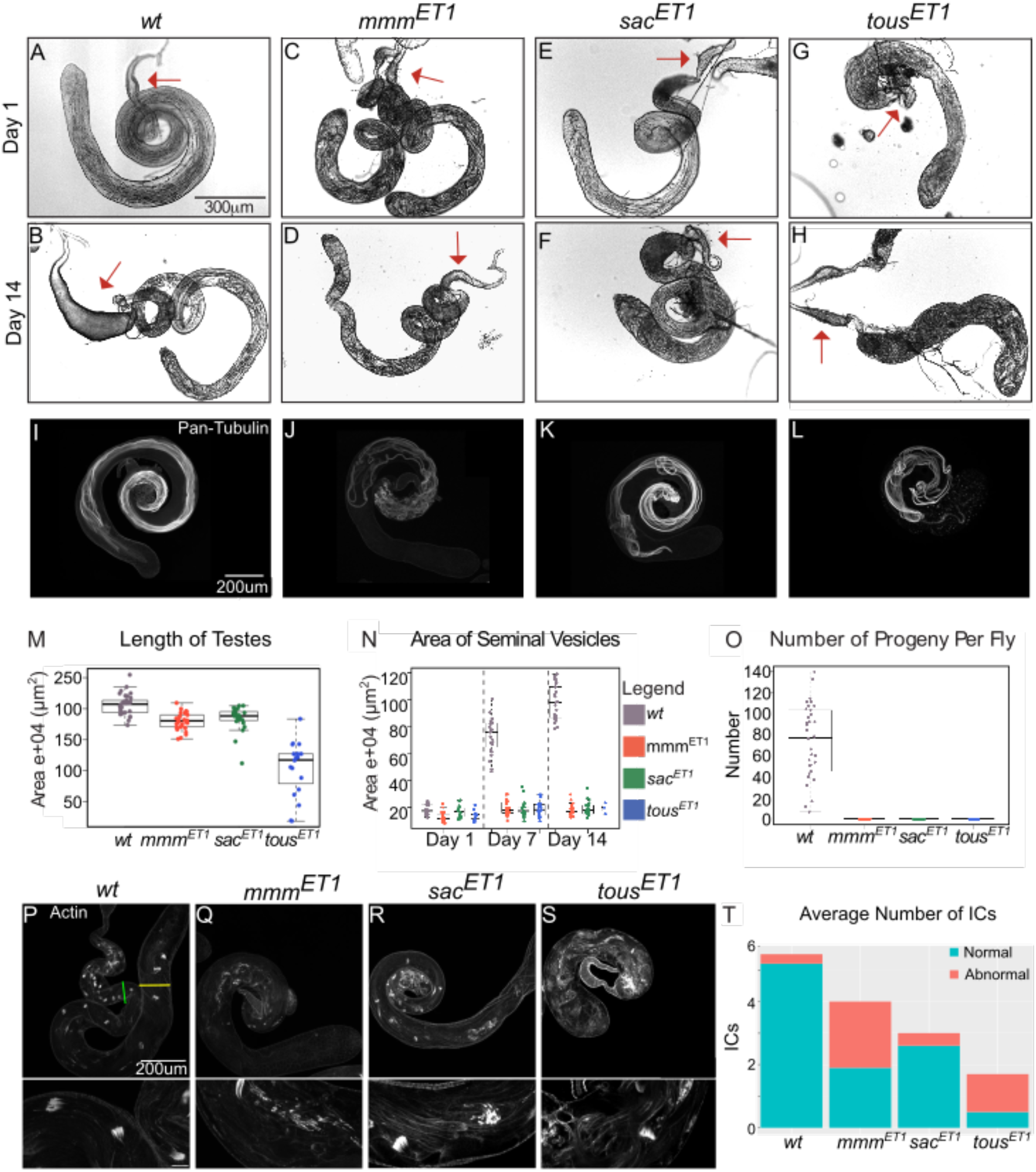
*mmm, sac*, and *tous* are required for spermatogenesis. (A-H) Brightfield images of live testes and associated seminal vesicles (*red* arrow) examined for morphology from (A-B) *wildtype,* (C-D) *mmm^ET1^*, (E-F) *sac^ET1^*, and (G-H) *tous^ET1^* homozygous males at days 1 and 14. (I-L) Whole mount testes labeled with Anti-pan polyglycylated Tubulin displayed mature sperm in (I) *wildtype (n*=14), (J) *mmm^ET1^* (*n*=14), (K) *sac^ET1^* (*n*=9), and (L) *tous^ET1^* (*n*=10) homozygotes. (M) Testis length of *wildtype (grey, n*=33), *mmm^ET1^ (red, n*=30), *sac^ET1^ (green, n*=23), and *tous^ET1^* homozygotes (*blue, n*=19). (N) Quantification of the seminal vesicle area from *wildtype (grey, n*=22 on day 1, *n*=24 on day 7, *and n*=25 on day 14), *mmm^ET1^ (red, n*=29 on day 1, *n*=23 on day 7, *and n*=24 on day 14), *sac^ET1^ (green, n*=16 on day 1, *n*=22 on day 7, *and n=21* on day 14), and *tous^ET1^* (*blue, n=*11 on day 1, *n=21* on day 7, *and n=*5 on day 14) un-mated homozygous males. From day 1 to14, *wildtype* males showed a steady increase in seminal vesicle area as mature sperm was loaded. The area of the seminal vesicles of all mutants remained unchanged throughout 14 days. (O) Progeny counts of the total number of progenies from *wildtype (grey, n*=34 individual males), *mmm^ET1^ (red, n*=30 individual males), *sac^ET1^ (green, n*=27 individual males), and *tous^ET1^* homozygotes (*blue, n*=33 individual males). (P-S) Whole mount testes labeled with SiR-Actin to visualize the individualization complexes in *wildtype* (P), *mmm^ET1^* (Q), *sac ^ET1^* (R), and *tous ^ET1^* homozygotes (S). The *green* line indicates the initiation site at the nuclei and the *yellow* line indicates the termination. (T) Quantification of the average number of individualization complexes (ICs) that formed normally (*blue*) or abnormally (*pink*) per testis. *wildtype n*=17, *mmm^ET1/ET1^ n*=14, *sac^ET1/ET1^ n*=20, and *tous ^ET/ET1 1^ n*=12.

Separation of sperm bundles prior to the completion of spermatogenesis is dependent on Actin-mediated resolution of the cytoplasmic bridges between sperm. At the end of the remodeling events during spermiogenesis, actin cones form at the nucleus in the basal end of the testes for each of the 64 sperm within a bundle. These cones subsequently travel along the entire length of the sperm, resolving the cytoplasmic bridges that connect them. To investigate potential requirements for *mmm*, *sac*, and *tous* in this process, we used immunofluorescence to examine the formation of actin cones in each mutant background. In *wildtype* testes, an average of 5-6 actin cones were typically observed between the formation site at the nucleus and the termination sites at the apical end of the testes (Figure 1P-T). *mmm*, *sac*, and *tous* mutants all contained fewer actin cones than wildtype, indicating errors in either forming or maintaining these structures. *mmm* and *tous* mutants also displayed an increased number of abnormal actin cones where the actin cones appeared fragmented, supporting the interpretation that they are unable to properly form or maintain these structures. Taken together, these results confirm that *mmm*, *sac*, and *tous* are essential for male fertility.

### *mmm, sac*, and *tous* are required at multiple stages of spermatogenesis

Nuclei and mitochondria undergo a series of major remodeling events during spermatogenesis, the first phase of which is dedicated to cell growth and division (Figure 2A). Spermatogonia mitotically divide to produce 16 spermatocytes that, after a prolonged growth phase, undergo meiosis resulting in 64 spermatids. During the next phase, spermatids enter spermiogenesis as mitochondria coalesce during the aggregation stage while ciliogenesis initiates. These events are immediately followed by the onion stage and elongation, in which the terminal differentiation of spermatids initiates. To visualize these stages in each mutant, we crossed *mmm* and *tous* mutant alleles with a *His2AV-mRFP1, sqh-EYFP-mito* transgenic line to recombine on fluorescent markers for DNA and mitochondria. The *sac* mutant was crossed with a *His2AV-mRFP1* transgenic line to recombine on a fluorescently labeled DNA marker and MitoTracker was used to visualize the mitochondria. Using phase-contrast microscopy in conjunction with epifluorescence, we examined spermatogenesis to determine which stages, if any, were perturbed. Prior to meiosis, cells from all three lines appeared morphologically indistinct from *wildtype* cells. As with *wildtype*, both *sac^ET1^* and *tous^ET1^* meiotic cells displayed DNA at either end of the spindle poles and mitochondria evenly distributed along the meiotic spindle (Figure 2B-C). However, 14% of *tous^ET1^* cells displayed abnormal localization of mitochondria along the meiotic spindle. Immediately after meiosis, all three mutant alleles displayed an increase in the number of cells undergoing mitochondrial aggregation. This stage is normally short-lived (30 minutes) and only one 64 cell cyst should be in this stage at any given time.As such, this stage is infrequently observed in wildtype, compared to stages such as the pre-meiotic S phase which lasts 4 days. However, in testes from all three mutants aggregating cells were readily observable compared to *wildtype* control, indicating a likely delay in the transition from meiosis into spermiogenesis (Figure 2D-H) (Fuller, 1993; Lindsley & Tokuyasu, K.T., 1980). Especially, *sac^ET1^* and *tous^ET1^* testes displayed a dramatic increase in the number of cells undergoing mitochondrial aggregation (Figure 2D). Consistent with this, evidence of impaired meiotic cytokinesis was observed in *tous^ET1^*, where nuclei of variable sizes at the onion stage were observed, a phenomenon caused by unequal distribution of DNA during cytokinesis (Figure 2L and M) (Castrillon et al., 1993, 1993; Fuller et al., 1988). While *mmm^ET1^* and *sac^ET1^* displayed less variability in size, both displayed enlarged nuclei. While the cause remains unclear, it could be due to a delay in nuclear condensation, which would be consistent with the delayed exit from the aggregation stage (Figure 2D-M). During the elongation stage, *wildtype* mitochondria typically form compact elongated tails that encapsulate the cilium, however, both *mmm^ET1^* and *tous^ET1^* displayed a loss of mitochondrial integrity (Figure 2N-Q). Both exhibited similar phenotypes characterized by an apparent disassembly of the mitochondrial derivate. *sac^ET1^* mutants were commonly observed in the earlier and less compact stages of elongation, appeared to be able to achieve normal elongation status. Collectively, our results suggest that *mmm*, *sac*, and *tous* are all required during meiosis and that *mmm* and *tous* are involved in mitochondrial integrity during spermiogenesis.

**Figure 2.**
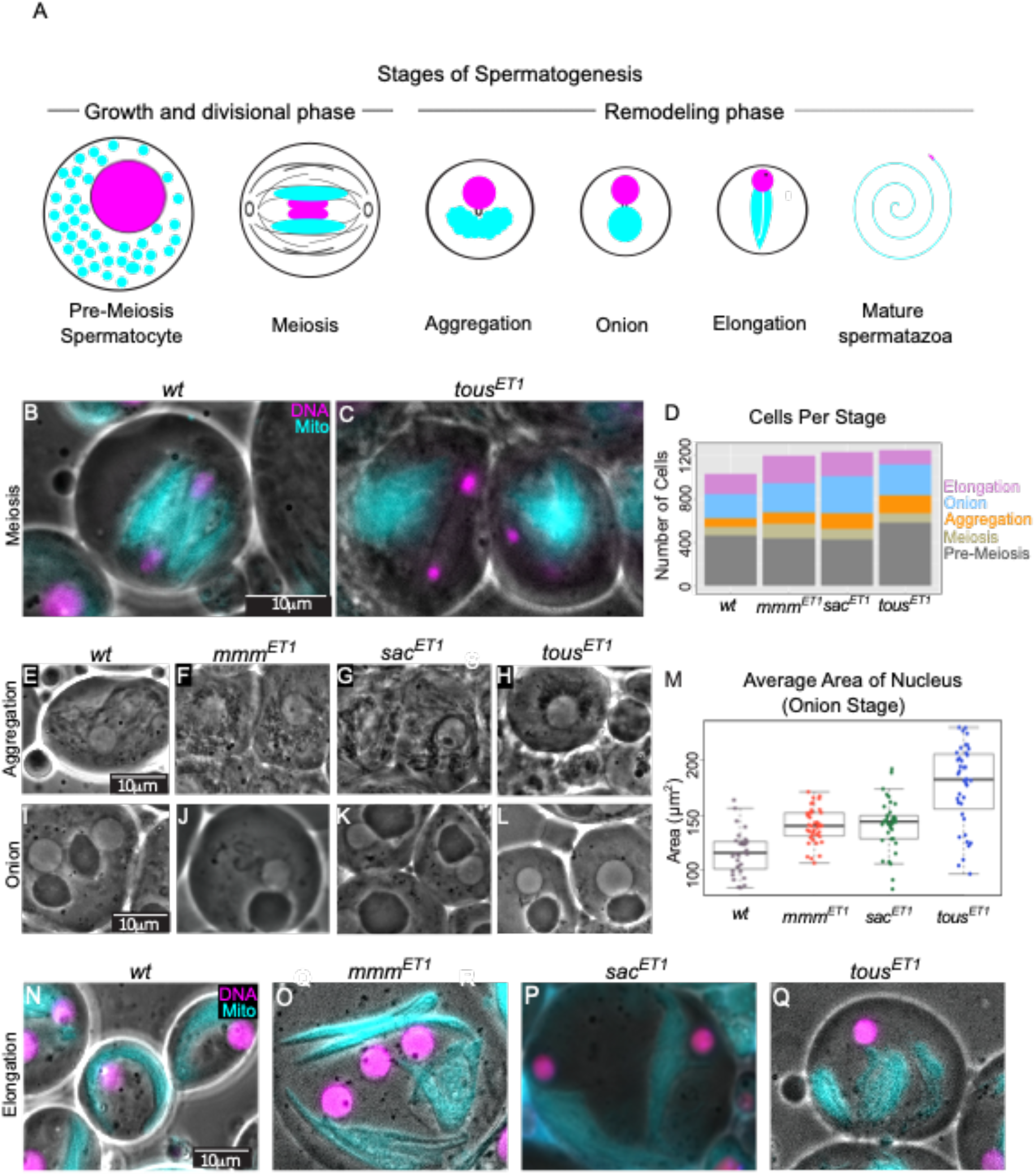
Phenotypes of *mmm, sac*, and *tous* mutants during spermatogenesis. (A) Schematic of cell stages throughout spermatogenesis (adapted from Bauerly et al., 2020). Prior to meiosis, cells are in a growth and divisional phase, increasing in size by ~25x and dividing from a single spermatogonium to 16 spermatocytes by the completion of the pre-meiotic S-phase. After the two rounds of meiosis, the resulting 64 spermatids enter a remodeling phase where the nucleus and mitochondria undergo drastic transformation concurrently with the initiation of ciliogenesis. (B-C) Phase-contrast microscopy with epifluorescence of spermatids during meiosis for *wildtype* (B) and *tous ^ET1^* (C) homozygous spermatids. Misaligned mitochondria were observed in 14% of *tous ^ET1 /ET1^* cells along the meiotic spindle (*n*=12/88). (D) Quantification of the number of cells during pre-meiosis (*brown, wildtype n*=465/1030 cells, *mmm^ET1/ET1^ n*=439/1197 cells, *sac^ET1/ET1^ n*=428/1232 cells, *tous^ET1/ET1^ n*=581/1249 cells), meiosis (*beige, wildtype n*=78/1030 cells, *mmm^ET1/ET1^ n*=132/1197 cells, *sac^ET1/ET1^ n*=97/1232 cells, *tous^ET1/ET1^ n*=88/1249 cells), aggregation, (*orange*, *wildtype n*=77/1030 cells, *mmm^ET1/ET1^ n*=105/1197 cells, *sac ^ET1/ET1^ n*=143/1232 cells, *tous^ET1/ET1^ n*=166/1249 cells), onion (*blue, wildtype n*=227/1030 cells, *mmm^ET1/ET1^ n*=271/1197 cells, *sac^ET1/ET1^ n*=341/1232 cells, *tous^ET1/ET1^ n*=281/1249 cells), and elongation (*pink, wildtype n*=183/1030 cells, *mmm^ET1 /ET1^ n*=250/1197 cells, *sac^ET1/ET1^ n*=223/1232 cells, *tous^ET1/ET1^ n*=133/1249 cells) stages. (E-L) Phase-contrast microscopy of live spermatids during the aggregation and onion stages for *wildtype* (E and I), for *mmm^ET1^* (F and J), *sac ^ET1^* (G and K), and *tous ^ET1^* (H and L) homozygotes. The increase in aggregation cells noted in (D) is readily observable in phase-contrast microscopy of live spermatids displaying cells compared to *wildtype* (E) for *mmm^ET1^* (F), *sac ^ET1^* (G), and *tous ^ET1^* (H) homozygotes. (M) Quantification of the size of the nucleus during the onion stage for *wildtype (n*=39*), mmm^ET1^(n*=31), *sac^ET1^(n*=37), and *tous^ET1^(n*=39) homozygotes. (N-Q) Phase-contrast microscopy with epi-fluorescence of live spermatids during elongation for *wildtype* (N, *n*=2/183), *mmm^ET1^* (O, *n*=30/250), *sac^ET1^* (P, *n*=7/223), and *tous^ET1^* (Q, *n*=28/133) homozygotes. Aberrant mitochondrial conformation can be observed in 12% of *mmm ^ET1/ET1^* and 21% of *tous ^ET1/ET1^* cells.

### *mmm, sac*, and *tous* are essential for the proper formation of cellular substructures

During the elongation phase, *wildtype* spermatids are encompassed by a plasma membrane which contains the developing axoneme flanked by the major and minor mitochondrial derivatives. The elongation of these two derivatives is in turn required for elongation and function of the flagellum (Noguchi et al., 2011; Tates, A.D., 1971). To investigate the cause of sperm immotility in *mmm^ET1^*, *sac^ET1^*, and *tous^ET1^* homozygotes, we next analyzed the ultrastructure of the developing axoneme using electron microscopy. While no obvious defects of the axoneme were observable, all three mutant alleles displayed defects of various cellular structures (Figure 3). In *mmm^ET1^* homozygotes, the major mitochondrial derivative appeared morphologically normal, however several cells lacked a corresponding minor mitochondrial derivative (Figure 3B). In *sac ^ET1^* homozygotes, the plasma membrane was abnormal in many cells, resulting in the commingling of numerous axonemes and mitochondria (Figure 3C). Similarly, the plasma membrane appeared abnormal in *tous^ET1^* homozygotes, bringing multiple axonemes and mitochondria into close proximity (Figure 3D). This phenotype has previously been described in cells with failed cytokinesis (Carmena et al., 1998) Additionally, in *tous* mutants, we observed that axonemes frequently either failed to form or lost the ability to maintain a mitochondrial derivative, resulting in multiple axonemes sharing a mitochondrial derivative (Figure 3D). Collectively, these data indicate a role for *mmm*, *sac,* and *tous* in either forming or maintaining the integrity of cellular structures during spermatogenesis.

**Figure 3.**
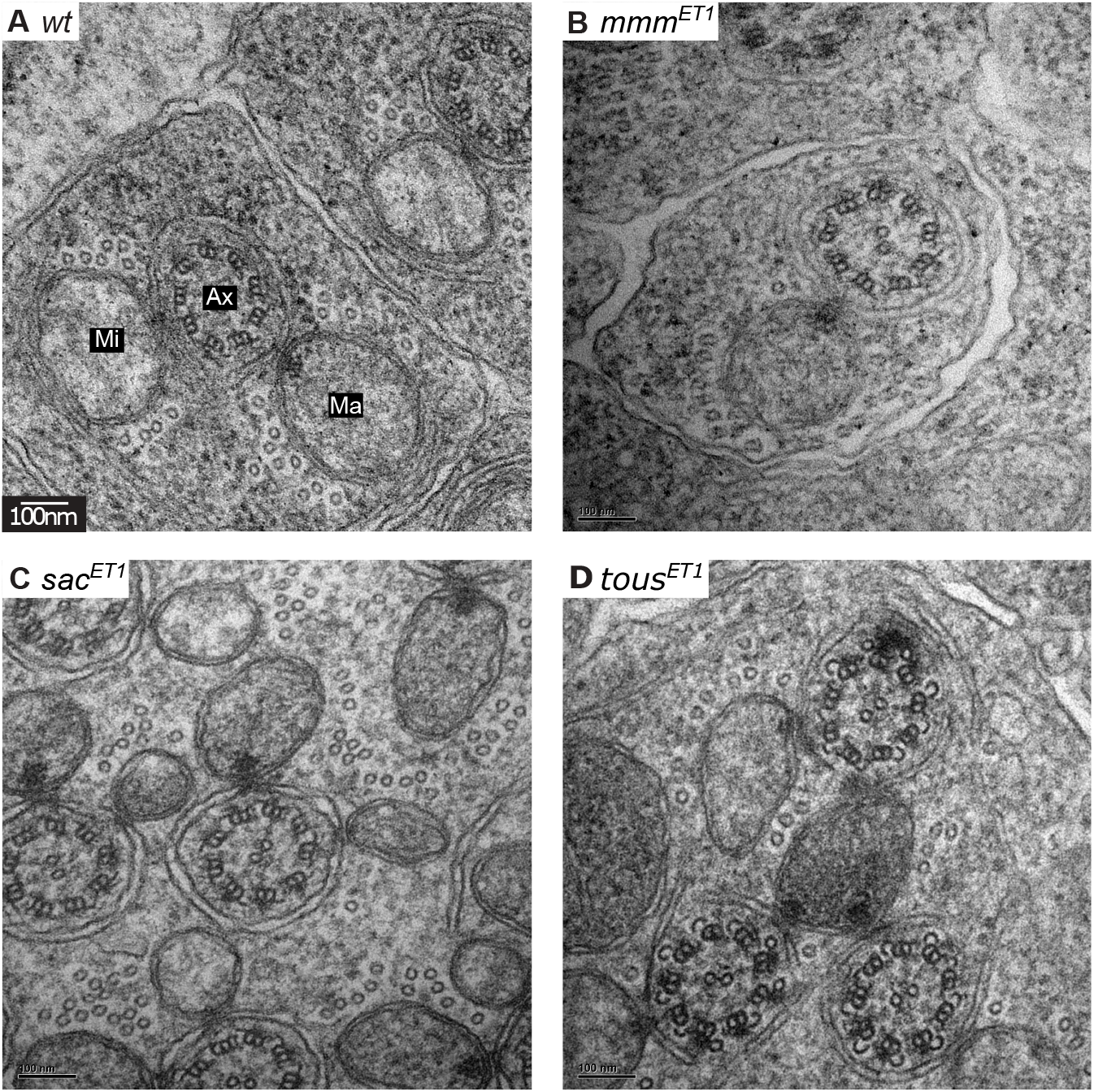
*mmm, sac*, and *tous* are essential for the proper formation of cellular substructures. (A-D) EM analysis of elongating spermatids showed the expected localization of the axoneme (Ax) relative to the major (Ma) and minor (Mi) mitochondrial derivatives, encompassed by the plasma membrane in *wildtype* (A, *n*=22). *mmm^ET1^* homozygous spermatids lack a minor mitochondrial derivative in a subset of cells (B, *n*=7/13). Both *sac ^ET1^* and *tous ^ET1^* homozygous spermatids displayed a loss of integrity of the plasma membrane (C and D; *sac ^ET1^ n*= 5/10, *tous ^ET1^ n*=10/16). *tous ^ET1^* homozygous spermatids also have one axoneme sharing multiple major mitochondrial derivatives (D, *n=* 3/16).

### Mitochondrial localization is aberrant in *mmm, sac*, and *tous* homozygous testes

While analyzing the subcellular organization developing spermatids, we observed that *wildtype* mitochondrial derivatives tend to display comparable sizes at equivalent developmental time points, whereas the size of the mitochondrial derivatives varied widely in the mutant backgrounds (Figure 4A-G). Intriguingly, the minor derivative appears to be particularly susceptible to perturbation, displaying a significant size reduction compared to *wildtype* in all three mutants (Figure 4G). The accessory microtubules that surround the mitochondrial derivatives are essential in the elongation process of the mitochondria (Hoyle & Raff, 1990). Given the elongation defects of the mitochondrial derivatives observed in these mutants, we next examined the accessory microtubules for defects that might explain the disruption to the development of the mitochondrial derivatives.

**Figure 4.**
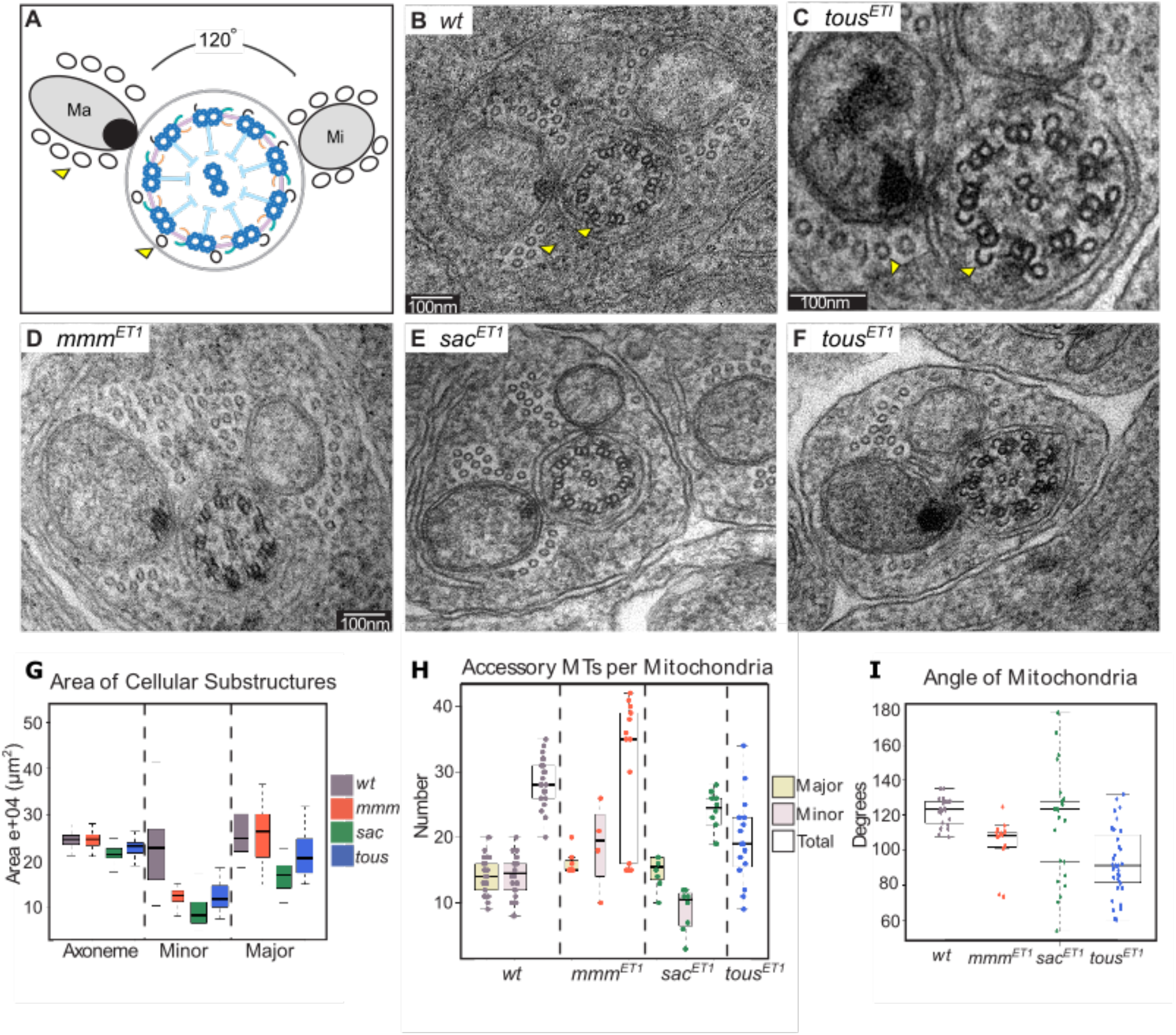
Mitochondrial formation is perturbed in *mmm, sac*, and *tous*. (A-F) (A) Schematic and (B) *wildtype* EM image displaying the axoneme (Ax) of an elongating spermatid with the major (Ma) and minor (Mi) mitochondrial derivatives approximately 120° degrees apart with respect to the central pair. (C) EM image of the axoneme of an elongating spermatid from *tous ^ET1/ET1^* revealing the formation of accessory microtubules (*yellow* arrow). EM images displaying the axonemes of elongating spermatids from (D) *mmm^ET1^*, (E) *sac^ET1^*, and (F) *tous^ET1^* homozygotes. The angle between the mitochondrial derivatives in (D) *mmm^ET1^* and (F) *tous ^ET1^* homozygous spermatids was significantly decreased, while the angle between the mitochondrial derivatives (E) *sac^ET1^* homozygous spermatids remained at the expected 120°, with more variation. (G) Quantification of the area of the major and minor mitochondrial derivatives and axonemes for *wildtype (grey, n*=26*), mmm^ET1^(red, n*=19), *sac^ET1^(green, n*=23), and *tous^ET1^ (blue, n*=27) homozygous spermatids. The axoneme was approximately the same size for all genotypes, but the minor mitochondrial derivatives in all three mutants showed a significant decrease in size compared to *wildtype*. Additionally, *sac ^ET1^* and *tous^ET1^* homozygous spermatids also displayed a notable reduction in the size of the major mitochondrial derivatives. (H) Quantification of the number of accessory microtubules per mitochondrial derivative. *wildtype* (*grey, n*=22) spermatids had an average of 14 accessory microtubules per mitochondria with an average of 28 per cell. *mmm^ET1^* (*red, n*=13) homozygous spermatids contained the expected number of accessory proteins surrounding the major mitochondrial derivative, but cells that also contained a minor derivate displayed an increase in the number of accessory microtubules, bringing the total to an average of 35 accessory microtubules per cell. *sac ^ET1^* homozygotes *(green, n*=10) contained the expected number of accessory microtubules around the major mitochondrial derivative, but had fewer surrounding the minor derivative, resulting in a lower total number of accessory microtubules. While the angle between the mitochondrial derivatives was too small to adequately distinguish between the accessory microtubules of the major and the minor derivative in *tous^ET1^*(*blue, n*=16) homozygous spermatids, the total number was an average of 21 per cell, slightly reduced from that of *wildtype.* (I) Quantification of the angle between the major and minor mitochondrial derivatives relative to the central pair. The angle of the mitochondria relative to the central pair averaged 120° in *wildtype (grey, n*=25) spermatids. However, this angle was reduced to 112° and 90° in *mmm^ET1^ (red, n*=15) and *tous^ET1^ (blue, n*=26) homozygous spermatids, respectively. While the angle averages the expected 120° in *sac ^ET1^ (green, n*=36) homozygous spermatids, there was increased variability relative to *wildtype.*

Accessory microtubules are generated from the β-tubule of the axoneme during the elongation stage (Tokuyasu, 1974) (*yellow* arrows, Figure 4A-C). After forming, they are released from the membrane, find their way to the mitochondria, and then divide in approximately equal proportions to encompass both derivatives. To address whether the accessory microtubules were affected in each mutant, we analyzed the distribution of accessory microtubules around both mitochondrial derivatives (Figure 4H). In *wildtype* cells, each derivative had an average of 14 associated microtubules surrounding it, resulting in approximately 28 total accessory microtubules per cell (Figure 4H). For *mmm^ET1^* homozygous testes, the cells that lacked a minor mitochondrion only had the approximately 14 accessory microtubules surrounding the major mitochondria. However, the cells that contained both derivatives seemed to have an increase in the number of accessory microtubules, resulting in an overall higher average than what is typically observed (Figure 4H). This suggests that the correct number of accessory microtubules might be generated and then distributed through the shared cytoplasm to neighboring minor mitochondrial derivates and implies the potential existence of a recognition mechanism that can distinguish between the major and minor derivatives, which would also account for the nearly equal distribution of accessory microtubules that is observed in *wildtype* cells.

*sac^ET1^* homozygotes, which had the strongest size reduction of the minor mitochondria, had fewer accessory microtubules associated with the minor derivative, resulting in a decrease in the total number of accessory microtubules per cell (Figure 4G-H). *tous ^ET1^* homozygous testes displayed a decrease in the total number of accessory microtubules per cell. However, due to the close proximity of the major and minor mitochondria, it was not possible to distinguish which microtubules were associated with a specific derivative (Figure 4D and H). Interestingly, *mmm^ET1^* cells had, on average, a higher total number of accessory microtubules and the major mitochondrial derivate size was consistent with *wildtype*. The size defect observed in the minor mitochondria in *mmm^ET^* cells is presumed to be related to a defect in either forming or maintaining this derivative. However, cells from both *sac^ET1^* and *tous ^ET1^* contained, on average, fewer accessory microtubules and both mitochondrial derivatives were smaller than what is observed in *wildtype* cells. This data suggests that there is a correlation between fewer accessory microtubules and smaller mitochondrial derivatives.

In addition to the irregularities in size, we also noted that the angle at which the derivatives contact the axoneme was strikingly altered in both *mmm^ET1^*and *tous^ET1^* homozygous testes (Figure 4A-F and I). In *wildtype* testes, the angle by which the derivatives contact the axoneme, relative to the central pair, is approximately 120° with little variation (Tokuyasu, 1974) (Figure 4I). This angle was significantly reduced in both *mmm ^ET1^*and *sac ^ET1^* homozygotes, obscuring the association of the accessory microtubules with the mitochondria in many cells. While *sac^ET1^* homozygotes displayed an average of 120° degrees between the mitochondrial derivatives, this angle was highly variable from cell to cell. It is not known what regulates the association of the mitochondria and the axoneme or the formation and localization of the accessory microtubules, however these results suggest that a correlation between the number of accessory microtubules and the formation of the mitochondrial derivatives.

## Discussion

During spermatogenesis, ciliogenesis is tightly coupled to the development of the cell body and it is becoming increasingly evident that defects that affect ciliogenesis hold potential to also disrupt cell division (Hua & Ferland, 2018; Bauerly et al., 2020). Here, we report three novel cilia-related genes, *mmm*, *sac*, and *tous,* required for spermatogenesis all of which are essential for male fertility and present with immotile flagella upon inspection (Figure 1 and Supplemental Movie 1). These genes also show evidence for roles during meiosis or immediately thereafter, resulting in a prolonged entry into spermiogenesis following meiosis and aberrant formation of the mitochondrial derivatives (Figure 2–4).

During the elongation phase, we observed that *mmm^ET1^* cells frequently lacked a minor mitochondrial derivative (Figure 3B). Interestingly, the cells that lacked a minor mitochondrial derivative only contained an average of 14 accessory microtubules and not the total average of 28 that was observed in cells with both derivatives (Figure 4H). The other 14 accessory microtubules that typically surround the minor mitochondria were not present in cells without a minor mitochondrion, suggesting that there is some feedback between the axonemal-derived accessory microtubules and the formation of the mitochondria. Consistent with this observation, mutant animals for both *sac* and *tous* had smaller minor mitochondrial derivatives and fewer associated accessory microtubules, suggesting a correlation between the formation of the mitochondria and the accessory microtubules (Figure 4H). While the formation and maintenance of mitochondrial derivatives are not well understood and the mechanisms regulating accessory microtubules have not yet been identified, it is intriguing that *mmm*, *sac*, and *tous* mutant animals all displayed size defects of the minor mitochondrial derivative and positioning defects for both mitochondrial derivatives relative to the axoneme. In *wildtype* testes, the mitochondria assume a fixed position relative to the axoneme throughout spermatogenesis (Tokuyasu, 1975). While dyneins are known to be involved in the regulation of mitochondrial dynamics, the mechanism surrounding their interaction is not completely understood (Fuller, Caulton, Hutchens, Kaufman, & Raff, 1987; Sitaram, Anderson, Jodoin, Lee, & Lee, 2012). The cumulative defects observed in the mitochondrial derivatives, particularly in the minor mitochondria, suggest that the minor derivatives are more sensitive to spermatogenic defects and that there is an unappreciated connection between the mitochondria and the accessory microtubules.

In summary, the present study identified and characterized cellular requirements for three novel genes with essential roles in spermatogenesis. These genes are necessary for ciliary function and display a range of defects throughout multiple stages of cellular development when disrupted. As the list of cilia-relate genes that function outside the cilium continues to grow, it is becoming increasingly clear that understanding how these proteins affect other aspects of cellular development will be indispensable for understanding the pleiotropic manifestation of diseases, such as sterility, that arise from the dysfunction of cilia.

## Supporting information

Supplemental Movie 1B_mmm

Supplemental Movie 1C_sac

Supplemental Movie 1D_tous

## Acknowledgements

We thank the Bloomington Stock Center for fly stocks, Steve Hoffman for assisting with microscopy, and members of the Avasthi lab for critical reading of the manuscript. This work was supported by funding from the Stowers Institute for Medical Research and NIH Grant R01-GM111733 to M.G..

## Author contributions

M.G. and E.B. designed the project. E.B. performed most experiments and data analysis. T.A. set crosses for and analyzed rescue data, K.Y. acquired EM images. M.G. and E.B. wrote the manuscript.

## Declaration of Interests

No competing interests

## Materials and Methods

### *Drosophila melanogaster* stocks

All stocks were maintained at 25°C on corn syrup food. The *wildtype* stock used was *w^1118^,* except when noted. Stocks for *mmm^ET1^* (BDSC#24682), *sac ^ET1^* (BDSC#27860), and *tous ^ET1^* (BDSC#23200) were obtained from the Bloomington *Drosophila* Stock Center.

The fluorescent lines used for imaging the mitochondria and DNA were created by crossing *His2Av-mRFP1,* a histone variant fused to RFP (BDSC#23650 and 23651) to *sqh-EYFP-Mito 3*, which contains a ubiquitously expressed mitochondrial targeting sequence under the control of *sqh* regulatory sequences to drive the expression of the EYFP (BDSC#7194), to generate the control lines *w; His2Av-mRFP1 II.2*; *sqh-EYFP-Mito* and *w; His2Av-mRFP1 III.1*, *sqh-EYFP-Mito 3*.

*mmm^ET1^* and *sqh-EYFP-Mito* were crossed to allow for recombination between the mitochondrial marker and the mutation onto the same chromosome. This line was then crossed with *His2Av-mRFP1 II.2* to generate *w; His2Av-mRFP1 II.2; mmm^ET1^, sqh-EYFP-Mito 3*. Due to the proximity of the *sac ^ET1^* mutation and *sqh-EYFP-Mito* marker, a recombinant could not be recovered. This line was instead crossed to *His2Av-mRFP1 II.2* to generate *w; His2Av-mRFP1 II.2; sac^ET1^*. MitoTracker green (Invitrogen, M7514) was used to label the mitochondria for this line and *His2Av-mRFP1 II.2* was used as the control. *His2Av-mRFP1 III.1* and *sqh-EYFP-Mito* were crossed to allow for recombination to generate *w; His2Av-mRFP1 III.1*, *sqh-EYFP-Mito*. *tous ^ET1^/*/; *Sp/TM3* was then crossed on to this line to create *w; tous ^ET1^//; His2Av-mRFP1 III.1*, *sqh-EYFP-Mito.*

The mutant lines were validated both by crossing to a deficiency line and to a rescue construct. The stocks used for verification were BDSC#30580, 7620, 282 for *mmm^ET1^*, *sac ^ET1^*, and *tous ^ET1^*, respectively, from the Bloomington *Drosophila* Stock Center. Both *mmm^ET1^* and *sac ^ET1^* recapitulated the sterile phenotype when crossed to a deficiency line. However, *tous ^ET1^* resulted in lethality when crossed to the deficiency line indicating that this line is either a hypomorphic allele or that the combination of the *tous ^ET1^* mutation with the disruption of another gene in the deficiency line is toxic. The rescue constructs were generated from a fragment encompassing the locus of interest plus 2 kb on either side. An mScarlet-I tag was placed at the C-terminus with a flexible 10x glycine linker that was incorporated into the primers. The following primers were used (5’-3’):

### CH322-31D12 (mmm)

5’ homology arm F: CCTGCAGGTCGACTCTAGAGGTTTAAACTCGCGATTATGATGGCAATG

5’ homology arm R: ACCGCCTCCTCCACCTCCGCCACCACCACCTCCGCCACCACCACCACCC TCGAAATCGTCCAC

3’ homology arm F: TAAAGTTTGTAATGCGTCAG

3’ homology arm R: ATTCGAGCTCGGTACCCGGGGTTTAAACAACTCTTTCATTTACTTTAT

### pmScarlet-i_C1

Scarlet-I F: GACGATTTCGAGGGTGGTGGTGGTGGCGGAGGTGGTGGTGGCGG AGGTGGAGGAGGCGGTATGGTGAGCAAGGGCGAGGC

Scarlet-I R: CTGACGCATTACAAACTTTACTTGTACAGCTCGTCCATGC

### CH322-178G16 (*sac*)

5’ homology arm F: CCTGCAGGTCGACTCTAGAGGTTTAAACATGCCGATGGCTATGGCGATGG

5’ homology arm R: ACCGCCTCCTCCACCTCCGCCACCACCACCTCCGCCACCACCACCTAGTATATCCCGGCGACACT

3’ homology arm F: TAGTTTGTTGTTGAAATCGT

3’ homology arm R: ATTCGAGCTCGGTACCCGGGGTTTAAACGCGATGGACAAAAATTTTA CAAGAAACACAAC

### pmScarlet-i_C1

Scarlet-I F: CGCCGGGATATACTAGGTGGTGGTGGCGGAGGTGGTGGTGGCGGAGGTGGAGG AGGCGGTATGGTGAGCAAGGGCGAGGC

Scarlet-I R: ACGATTTCAACAACAAACTACTTGTACAGCTCGTCCATGC

### CH322-9O8 (*tous*)

5’ homology arm F: CCTGCAGGTCGACTCTAGAGGTTTAAACTTGCTGGC TGACAGAGTCCA

5’ homology arm R: ACCGCCTCCTCCACCTCCGCCACCACCACCTCCGCC ACCACCACCCTTGTGTATTGTCTTTGTTG

3’ homology arm F: TAGACATGCGAGTCTATGTG

3’ homology arm R: ATTCGAGCTCGGTACCCGGGGTTTAAACCCAAGAA GGAGATCAAGGACA

### pmScarlet-i_C1

Scarlet-I F: AAGACAATACACAAGGGTGGTGGTGGCGGAGGTGGTGGTG GCGGAGGTGGAGGAGG CGGTATGGTGAGCAAGGGCGAGGC

Scarlet-I R: CACATAGACTCGCATGTCTACTTGTACAGCTCGTCCATGC

The fragments were gel extracted using Zymoclean Gel DNA Recovery kit (Zymo Research, D4007), assembled using HIFI DNA assembly master mix (New England BioLabs, Inc, E261S) for an hour at 50°C and transformed into electrocompetent cells. The insert was then cloned into the *w+attB* plasmid (addgene, 30326) by digesting with PmeI and gel purified. *w+attB* donor plasmid was digested with XhoI, blunted with Klenow fragment, treated with shrimp alkaline phosphatase, and purified. The insert and vector were then ligated, transformed, screened, and sequence verified. The DNA rescue constructs for *mmm* and *sac* were integrated into *VK1(2R)59D3* and the *tous* DNA construct into *CK22(3L)65B2* attP attachment site via phiC31integrase-mediated transgenesis (ref).

To test the functionality of the rescue constructs, they were double balanced on the second and third chromosomes and crossed to *mmm^ET1^*, *sac^ET1^*, and *tous^ET1^*. Homozygous males from this stock were collected and assayed for fertility. *mmm^ET1^*and *sac ^ET1^* were able to rescue in 9/10 crosses using individual males crossed to *w^1118^* females. The *tous* rescue construct was unable to rescue, this is believed to be due to the tag disrupting the function of the rescue. Consistently, this construct was unable to be maintained in a stock when crossed to the *tous ^ET1^* mutant stock.

### Fertility assays

For all fertility assays, a minimum of 30 males were singly crossed to 3-5 *wt* (*w^1118^*) virgins except where noted. Vials with progeny after 10 days were considered fertile.

### Live imaging

Sperm motility was monitored by dissecting testes from 2-3 day old males that were transferred to a clean slide in a drop of PBS (pH 7.4). The testes and seminal vesicles were pierced to release the sperm and imaged every 2ms for 100 frames on a Zeiss Axiovert 200M microscope using a 20x 0.8 Phase Plan-Apochromat objective.

Testes and seminal vesicle morphology were analyzed by collecting several newly eclosed homozygous males from *wt*, *mmm^ET1^*, *sac ^ET1^*, and *tous ^ET1^* and holding them as virgins for two weeks. Every few days males were analyzed. Their testes and associated seminal vesicles were dissected in PBS and transferred to a clean slide into a drop of PBS and imaged on a Leica CTR 5000 compound microscope.

Testes squashes were performed as previously described (Bauerly, et. al., 2020). In brief, testes were dissected from 0-2 day old homozygous males and transferred to a clean poly-L-lysine coated slide into a drop of PBS (pH 7.4). MitoTracker Green was used for *sac ^ET1^* and the control (see above) The testes were pierced 2-3 times to release the cells. The squashes were imaged on a Zeiss Axiovert 200M microscope using either a 20x 0.8 Phase Plan-Apochromat or 100x 1.4 Phase Plan-Apochromat objective. Brightness was occasionally adjusted for clarity and to avoid saturation.

### Fixed imaging

Whole testes from 0-2 day homozygous males were dissected in PBS (pH 7.4) and fixed in 4% paraformaldehyde for 25 minutes. Samples were then washed three times in 0.1% PBS+Triton x-100 (pH 7.4) for 15 minutes each. Hoechst was added at 1:1,000 and incubated overnight at 4°C. Samples were again washed three times in 0.1% PBS+Triton x-100 for 15 minutes each and then incubated in SiR-Actin, a highly specific fluorescent dye for F-actin (1:250, Cytoskeleton, Inc., CY-SC001), for 3 hours at room temperature. Samples were then mounted in Vectashield (Vector Laboratories, H-1000-10) and imaged on a Leica TCS SP5 confocal microscope.

### EM analysis

For TEM analysis, testes were dissected and fixed with a buffer containing 2.5% paraformaldehyde, 2% glutaraldehyde, 1% sucrose, and 50 mM sodium cacodylate (pH 7.4). After a brief rinse, the tissue was post-fixed with 1% OsO4 for 90 minutes and then stained with 1% uranium acetate en bloc overnight. Thereafter, the samples were dehydrated through an ethanol gradient to 100%, equilibrated with propylene oxide, and infiltrated with 50% propylene oxide/50% Epon, then 100% Epon resin (EMS, Fort Washington, PA) for 3 times over a day. After polymerizing at 60°C for 48 hours, the sample block was sectioned with a Leica Ultramicrotome (Leica UC-6) using diamond knives. The sections were post-stained with uranyl acetate and lead citrate, and then imaged with an FEI transmission electron microscope (Tecnai Bio-TWIN 12, FEI).

### Image analysis and quantification

All analysis was performed using ImageJ (Schneider, Rasband, & Eliceiri, 2012). Brightness was adjusted as needed to avoid saturation. Area quantifications were performed on images acquired on the same scope, with the same volume of media, with the same settings using the ‘Measure’ function. Statistical analyses were performed using a two-sample t-test, *n* values are listed in the figure legends.

## Notes

### Competing Interest Statement

The authors have declared no competing interest.

